# Autonomic/Central Coupling during Daytime Sleep Differs between Older and Younger People

**DOI:** 10.1101/2020.09.22.297184

**Authors:** Pin-Chun Chen, Katharine C. Simon, Negin Sattari, Lauren N. Whitehurst, Sara C. Mednick

## Abstract

Age-dependent functional changes are mirrored by declines in both the central the autonomic nervous systems and have been related to pathological aging. Prior studies in our group have identified a temporal coupling of Autonomic and Central Events (ACEs) during sleep using electrocardiogram to measure heart rate and electroencephalography to measure brain rhythms, with heart rate bursts (HRBs) temporally coincided with increased slow-wave-activity (SWA, 0.5-1Hz) and sigma activity (12-15Hz), followed by parasympathetic surge (RR_HF_) during non-rapid eye movement (NREM) sleep. ACEs predicted working memory (WM) improvement in young adults. Given that there are paralleling age-related declines in both the ANS and CNS, the current study investigated differences in ACE activity during daytime sleep in older and younger adults and their functional impact on working memory. Compared to youngers, older adults showed lower amplitude of ACEs during NREM sleep, but no age-related difference during Wake. Furthermore, while younger adults demonstrated a parasympathetic surge after HRBs, older adults showed a different pattern, with a earlier rise and maintenance of the RR_HF_. Finally, we examined whether ACE predicted WM in older adults. We found that older adults with good WM show stronger coupling, whereas low WM performers had less robust ACE activity. Taken together, our results demonstrated that autonomic-central coupling declines with age, with possible links to deterioration in WM function. Given that age-related deterioration in autonomic and central nervous system activity is implicated in pathological decline, the current findings may facilitate novel insights to the cognitive neuroscience of aging.

## 1. Introduction

Sleep supports a wide range of health and cognitive outcomes including restorative physiological processes, immune functioning, attention, learning and memory, creativity and semantic knowledge integration. The brain and body undergo large physiological changes across wake, NREM and REM that are thought to contribute to sleep’s benefits. REM sleep, is associated with faster, theta activity (4-7Hz) and phasic bursts of rapid eye movements visible in the electrooculogram (EOG). NREM sleep, on the other hand, is comprised of N2 and N3 and is characterized by a gradual slowing of the electroencephalagram (EEG) signal, including increases in slow waves that are critical for the facilitation of long-term memory formation (Marshall and Born, 2007; Diekelmann and Born, 2010; Rasch and Born, 2013). Slow wave activity (0.5-1Hz, SWA) is product of synchronous neuronal firing and a marker of the homeostatic regulation of sleep that has been linked to synaptic plasticity and cortical reorganization (Tononi and Cirelli, 2003; Takashima et al., 2006; Dang-Vu et al., 2010), as well as cerebral restoration and recovery (Horne, 1992).

In addition to EEG activity, sleep exerts considerable influence over the two branches of the ANS (i.e., sympathetic and parasympathetic nervous systems) with changes in the ANS modulating sleep onset and the transition between sleep stages (Trinder et al., 2001). As the brain shifts from wake into sleep, the body also undergoes marked changes with heart rate deceleration and relative increases in parasympathetic activity across NREM sleep. NREM sleep has been called a “cardiovascular holiday”, compared to wake and REM, due to it’s characteristic slower heart rate, reduced overall cardiac activity, and relative dominance of parasympathetic activity, and the ANS profiles during sleep has been viewed as an indication of cardiovascular health (Trinder et al., 2012). Although both CNS and ANS activity contribute to the benefits of sleep, little is known about how the two system interact and contribute to cognitive function.

Emergent literature has explored the association between central and autonomic activity during sleep, offering promising new perspectives to understand brain-body interplay in humans. de Zambotti et al (2016) investigated the relationship between cardiac activity and K-complexes (KC; 0.5-1 Hz)-a positive-negative-positive waveform during Stage 2 sleep similar to slow oscillations-and demonstrated that KCs were associated with a biphasic cardiac response, with a marked heart rate acceleration (de Zambotti et al., 2016). Interestingly, this biphasic fluctuation in heart rate has also been shown to coincide with bursts of K-complexes and delta waves (Sforza et al., 2000), which, together, indicate a synchronization of central and autonomic events. Additionally, ECG activity has been shown to modulate sleep spindle phase (Brandenberger et al., 2001; Lecci et al., 2017). More recently, our group evaluated changes in EEG and ECG signals that occurred during brief acceleration in heart rate during daytime nap (Naji et al., 2019). Naji et al. (2019) identified cardiovascular events during NREM sleep, termed heart rate bursts (HRBs), that last 2-3 seconds and occur mostly in Stage 2 and SWS (Naji et al., 2019). Heart rate bursts are temporally coincident with EEG activity, including SWA and spindle/sigma activity (12-15Hz) and are followed by vagal rebound reflected in a surge in the high frequency component (HF; 0.15–0.4 Hz) of the ECG. The increased SWA and Sigma activity before the HRBs and the vagal activity after the HR burst were termed as ACE activity (ACE-SWA, ACE-Sigma, and ACE-RRHF, respectively). Importantly, these Autonomic/Central Events (ACEs) positively predicted post-nap improvement in long-term declarative (Naji et al., 2019) and working memory (Chen et al., 2020). In fact, regression models showed a greater proportion of the sleep-dependent memory improvement was explained by the ACE activity than by the typical EEG sleep events alone.

With increasing age, sleep and associated cognitive functions decline (Mander et al., 2017). These age-dependent changes parallel dramatic alterations in sleeping brain rhythms between middle and late adulthood that are thought to reflect diminishing underlying neural integrity (Latreille et al., 2019; Mander et al., 2017). In older compared to younger adults, SWA shows reduced density (number of events/time) and amplitude in the prefrontal cortex (Carrier et al., 2011; Mander et al., 2013). In conjunction, sleep spindles show similar declines in density, length of burst, amplitude, and topography (Dijk et al., 1989; Mander et al., 2014; Martin et al., 2013). Declining sleep EEG physiology and general sleep characteristics, such as changes in sleep rhythms that include increased napping, nighttime arousal, and fragmented sleep, can prognostically predict future cognitive decline and are related to increased pathology and mortality (Dew et al., 2003; Keage et al., 2012).

Age-related declines in the ANS are also apparent (Hotta & Uchida, 2010). As we age, ANS functions can become more rigid, with decreased β-adrenoreceptor response, decreases in baroreflex activity, diminished heart rate responses to acetylcholine, and decreases in cardiac parasympathetic activity (Chadda et al., 2018). Studies show that these age-related ANS changes may contribute to autonomic imbalance, in which the sympathetic system is hyperactive and/or the parasympathetic system is hypoactive. Autonomic imbalance is associated with various pathological conditions, such as cardiovascular morbidity and mortality, diabetes, and dementia (Jandackova et al., 2016; Thayer et al., 2012; Villareal et al., 2002).

Given the independent lines of research showing age-related changes in CNS and ANS during sleep, research examining the interaction between ANS and CNS is necessary. The current study investigated whether aging impacts ACE activity during a daytime nap, as well as examining the functional impact of this activity on WM improvement over daytime sleep. 50 young and 51 older adults took a daytime nap in the lab with EEG and ECG monitoring. We predicted a reduction in ACEs for older compared to younger adults, such that the ACE-SWA and ACE-Sigma activity and ACE-RR_HF_ would be reduced in older compared to younger adults. Finally, we examined whether ACE predicted WM in older adults. We found that older adults with good WM show stronger coupling, whereas low WM performers had less robust ACEs. These results suggest that not only to the central and autonomic nervous systems individually change across the lifespan, but so does the strength of their interaction. Further research can investigate how this may impact cognition and other areas of functioning, which may help to identify novel areas for age-based interventions that may enhance quality of life.

## 2. Methods

### 2.1 Participants

Fifty healthy young adults (18-23yo, Mean =20.8, SD = 6.453) and fifty-one older adults (60-85yo, Mean = 69.99, SD = 3.074) with no personal history of neurological, psychological, or other chronic illness provided informed consent, which was approved by the University of California, Riverside Human Research Review Board. For the young adults, we reanalyzed a previously published data set (Chen et al., 2020). Participants included in the study had a regular sleep wake schedule (reporting a habitual time in bed of about 7–9 h for young adults and 6-8 h for older adults per night; National Sleep Foundation, 2015). Demographics and prior self-reported sleep habits were reported in Table 1.

**Table 1.** Descriptive statistics for Demographics

|  | *Young Adults (N=50)* | *Older Adults (N=51)* |  |
| --- | --- | --- | --- |
| Age (years) | 20.8 (6.453) | 69.99 (3.074) | *** |
| Male/female | 27/23 | 24/27 |  |
| Caffeine/day | 1.08 (1.554) | 1.82 (1.629) | *** |
| Alcohol/week | 0.47 (3.137) | 1.77 (3.311) | *** |
| Nap/week | 2.15 (2.635) | 2.07 (2.762) | *n.s.* |
| Nap duration/ day | 45.5 (52.386) | 36.8 (54.926) | *n.s.* |
| ESS | 7.39 (3.397) | 6.56 (4.619) | * |
| BMI (kg/m<sup>2</sup>) | 24.5 (6.706) | 27.7 (7.054) | *** |

*Note: Data are reported as Mean (standard deviation) for quantitative variables and N for categorical variables; ESS: Epworth Sleepiness Scale; BMI: Body Mass Index; Asterisks indicate significant differences between age groups (n.s. $p > 0.05$ ; \* $p < 0.05$ ; \*\*\* $p < 0.001$ ).*

The personal health histories were measured twice. First, during a pre-screening questionnaire in which a general health condition report was collected over the online survey and the eligibility was determined. Second, eligible subjects were invited to participate in an orientation in which they were given details about the study and interviewed by a trained graduate student. Participants who met any of the following exclusion criteria were excluded from the study: a) extreme morning or evening-type tendencies (measured with the Morningness Eveningness Questionnaire; Horne & Östberg, 1976); b) excessive daytime sleepiness (reported by Epworth Sleepiness Scale; Johns, 1991; subjects’ rating > 13 were excluded); c) a sleep disorder (assessed by questionnaires); d) any personal or immediate family (i.e., first degree relative) history of diagnosed significant psychopathology; e) personal history of head injury with loss of consciousness greater than 2 minutes or seizures; f) history of substance dependence; g) current use of any psychotropic medications; h) any cardiac, respiratory or other medical condition which may affect cerebral metabolism; i) non-correctable vision and auditory impairments.

Participants who did not met any of the exclusion criteria above and also met all the following inclusion criteria were enrolled in the study: a) aged 18-39/ 60-85 years old; b) healthy, non-smoking adult without major medical problems; c) completed at least 12 years of education; d) a regular sleep-wake schedule, defined as obtaining 7–9 h (young adults) or 6-8 h (older adults) of sleep per night, with a habitual bedtime between 9pm and 2am (young adults) or 8pm and 1am (older adults) and a habitual wake time between 6am and 10am (young adults) or 5am and 9am (older adults). Enrolled participants were asked to maintain their schedule for one week prior to their visit, which was monitored with sleep diaries. In addition, participants were asked to wear an actigraph (Actiwatch Spectrum, Respironics) for one night prior to their visit. Subjects were rescheduled if they reported poor sleep quality in their sleep diary, such as having more than 2 nights of less than 6 h of sleep or more than 9 h of sleep during the week prior to their visit, or if subjects’ actigraphy data showed less than 6 h of sleep or more than 9 h of sleep the night before the experimental visit. Rescheduled subjects were given another week to fill out a new sleep diary and maintain a regular sleep wake schedule prior to their visit.

During the orientation, participants were screened for cognitive impairment using Digit Span Backwards. Older participants were screened for dementia using the Telephone Screening for Dementia questionnaires (referred to as TELE), which consists of variety of questions to identify dementia-like symptoms. For older subjects, the STOP-BANG questionnaire was used to screen for obstructive sleep apnea and those with a medium to high risk were excluded. Additionally, participants were instructed to abstain from caffeine and alcohol starting at noon the day prior to the study (detected on the sleep diary). Participants who consumed alcoholic beverages > 10 cans a week or caffeinated products > 3 cups a day were excluded prior to the study (reported in Table 1).

### 2.2 Procedures

Participants were asked to maintain a regular sleep-wake schedule for one week prior to their visit, which was monitored with sleep diaries and was based on a 2-hour bedtime/ waketime window assigned at the orientation. The 2-hour bedtime/wake time was chosen based on subjects’ regular bedtime/wake time. In addition, participants were asked to wear an actigraphy (Actiwatch Spectrum, Respironics) for one night prior to their visit. Participants were rescheduled if they reported poor sleep quality in their sleep diary, such as having more than 2 nights of less than 6 h of sleep during the week prior to their visit, or if subjects’ actigraphy data was showing less than 6 h of sleep the night before the experimental visit. Rescheduled subjects were given another week to fill out a new sleep diary and maintain a regular sleep wake schedule prior to their visit.

On the experimental day, participants arrived at the Sleep and Cognition lab at 10:00AM and had EEG electrodes attached, followed by an Operation Span (OSpan) working memory task. Nap occurred between 1:30-3:30 PM. At 1:30PM, subjects were given 2-hourstime-in-bed to obtain up to 90-min total sleep time, monitored with polysomnography (PSG), including electroencephalography (EEG), electrocardiogram (ECG), electromyogram (EMG), and electrooculogram (EOG).. Sleep was monitored online by a trained sleep technician. All subjects were retested on the memory task between 4 and 4:30PM. Among our participants, 6 of older adults were not able to fall asleep, so they were excluded from further analyses. Participants received monetary compensation for participating in the study.

### 2.3. Data acquisition and pre-processing

#### 2.3.1 Sleep recording and scoring

Electroencephalographic (EEG) data were acquired using a 32-channel cap (EASYCAP GmbH) with Ag/AgCI electrodes placed according to the international 10-20 System (Jasper, 1958). Electrodes included 24 scalp, two electrocardiography (ECG), two electromyography (EMG), two electrooculography (EOG), 1 ground, and 1 on-line common reference channel. The EEG was recorded with a 1000 Hz sampling rate and was re-referenced to the contralateral mastoid (A1 & A2) post-recording. High pass filters were set at 0.3 Hz and low pass filters at 35 Hz for EEG and EOG. Only eight scalp electrodes (F3, F4, C3, C4, P3, P4, O1, O2), the EMG and EOG were used in the scoring of the nighttime sleep data. Sleep scoring were performed by two trained sleep technicians. Prior to scoring these data, technicians were required to reach 90% reliability with each other and one other rater on two independent data sets. Raw data were visually scored in 30-sec epochs into Wake, Stage 1 Sleep (N1), Stage 2 Sleep (N2), Slow Wave Sleep (N3) and rapid eye movement sleep (REM) according to the American Academy of Sleep Medicine (AASM) rules for sleep staging using HUME, a custom MATLAB toolbox. Prior to sleep scoring, data were pre-processed using BrainVision Analyzer 2.0 (BrainProducts, Munich Germany) and all epochs with artifacts and arousals were identified by visual inspection and rejected. Minutes in each sleep stage were calculated and sleep onset latency (SOL) were calculated as the number of minutes from lights out until the initial epoch of sleep, N2, N3 and REM. Additionally, wake after sleep onset (WASO) was calculated as total minutes awake after the initial epoch of sleep and sleep efficiency (SE) was computed as total time spent asleep after lights out divided by the total time spent in bed x 100. Descriptive statistics for sleep architecture were shown in Table 2.

**Table 2.** Summary of Sleep Architecture

|  | *Young* | *Older* |  |
| --- | --- | --- | --- |
| TIB (min) | 78.069 (3.1) | 80.763 (3.36) | *n.s.* |
| TST (min) | 61.500 (2.94) | 46.987 (3.92) | ** |
| SOL (min) | 7.664 (0.99) | 9.118 (1.27) | *n.s.* |
| N1 (min) | 4.586 (0.47) | 7.934 (1.07) | ** |
| N2 (min) | 31.448 (1.94) | 28.855 (2.86) | *n.s.* |
| N3 (min) | 19.224 (1.96) | 7.474 (1.58) | *** |
| REM (min) | 6.241 (1.09) | 2.72 (0.98) | * |
| WASO (min) | 6.560 (0.88) | 19.01 (2.12) | *** |
| SE (%) | 77.588 (2.21) | 56.52 (3.47) | *** |

*Note: Data are reported as Mean (standard error); TIB = Total Time in Bed; TST = Total Sleep Time; N3 = Slow Wave Sleep; REM = Rapid Eye Movement sleep; WASO = Wake After Sleep Onset (calculated as the minutes of wake after first epoch of sleep); SOL = Sleep Onset Latency (calculated as the time to first epoch of sleep); SE = Sleep Efficiency (calculated as $100 \times \text{TST} / \text{TIB}$ ). Asterisks indicate significant differences between age groups (n.s. $p > 0.05$ ; \* $p < 0.05$ ; \*\* $p < 0.01$ ; \*\*\* $p < 0.001$ ).*

#### 2.3.3 Power spectral analysis

The EEG power spectrum was computed using the Welch method (4 sec Hanning windows with 50 % overlap). The frequency for sigma power was 12-15Hz and for SWA was .5–1 Hz. For RR time-series, the power spectral estimation was performed by the autoregressive model and the model order was set at 16. Summary statistics for overall EEG power averaged across frontal and central areas (F3, F4, C3, and C4 channels) during each sleep stage were shown in Table 3.

**Table 3.** Summary of overall EEG Power Across Sleep Stages

| Stage | N2 |  |  | N3 |  |  | Wake |  |  |
| --- | --- | --- | --- | --- | --- | --- | --- | --- | --- |
|  | Young | Older |  | Young | Older |  | Young | Older |  |
| SWA (0.5-1Hz) | 89.785 (10.328) | 51.633 (5.658) | ** | 185.408 (14.037) | 124.643 (19.396) | * | 202.363 (24.726) | 107.083 (18.606) | * |
| Sigma (12-15Hz) | 5.30 (0.360) | 4.26 (0.442) | n.s. | 6.63 (0.724) | 4.07 (0.603) | * | 19.63 (3.387) | 10.12 (7.732) | n.s. |

Data are reported as Mean (standard error).
Asterisks indicate significant differences between age groups (n.s. $p > 0.05$ ; \* $p < 0.05$ ; \*\* $p < 0.01$ ; \*\*\* $p < 0.001$ ).

### 2.4 Autonomic/Central Event (ACE) Detection

For ACE analyses, consecutive artifact-free 3-min windows of undisturbed sleep (free from stage transitions, arousal, or movements) were selected across the whole nap. Epochs of N1 were not analyzed as only 2 older adults and 4 young adults had stable 3-min bins. For some subjects, we were unable to detect stable 3-min bins during other sleep stages, so they were excluded. In summary, we included 30 young and 42 older adults in analyses for Wake, 49 young and 42 older adults in analyses for N2, 38 young and 23 older adults in analyses for N3, as well as 19 young and 5 older adults in analyses for REM sleep. Since too few older subjects had REM sleep, we dropped REM from all the analyses.

#### 2.4.1 Heart-beat detection

Electrocardiogram (ECG) data were acquired at a 1000-Hz sampling rate using a modified Lead II Einthoven configuration. We analyzed HRV of the R-waves series using Kubios HRV Analysis Software 2.2 (Biosignal Analysis and Medical Imaging Group, University of Kuopio, Finland), according to the Task Force guidelines (Electrophysiology Task Force of the European Society of Cardiology the North American Society of Pacing, 1996). RR peaks were automatically detected by the Kubios software and visually examined by trained technicians. Incorrectly detected R-peaks were manually edited. Next, the data was passed through a lab tool based in MATLAB to perform further analyses. The ECG signals were filtered with a passband of 0.5-100 Hz by Butterworth filter. R waves were identified in the ECG using the Pan-Tompkins method, and confirmed with visual inspection. In order to extract continuous RR tachograms, the RR intervals were resampled (at 4 Hz for power spectrum estimation) and interpolated by piecewise cubic spline. Zero-phase Butterworth filters were applied to the interpolated RR time-series to extract RR_HF_.

#### 2.4.2 HR burst detection

Within each continuative and undisturbed 3-min bin, the mean and standard deviation of RR were calculated, and HR bursts were identified as the RR intervals shorter than 1.25 standard deviations below the mean. We investigated EEG changes during 20-sec windows around the HR bursts because EEG fluctuation typically returned to baseline level within 20 seconds in the previous study from our group (Naji et al., 2019). No other burst occurred in the 20 second windows around the HR bursts. Summary for HR bursts Density were shown in Table 4.

**Table 4.** Summary of HR Burst Density (per Minute) and RR intervals Across Sleep Stages

|  | N2 |  |  | N3 |  |  | Wake |  |  |
| --- | --- | --- | --- | --- | --- | --- | --- | --- | --- |
|  | Young | Older |  | Young | Older |  | Young | Older |  |
| HR Burst Density | 0.935 (0.0513) | 0.616 (0.0522) | *** | 0.892 (0.0563) | 0.557 (0.0673) | *** | 0.357 (0.0552) | 0.433 (0.0496) | n.s. |
| RR intervals | 991 (18.6) | 920 (22.1) | * | 988 (19.0) | 917 (23.5) | * | 920 (19.1) | 896 (22.5) | n.s. |

Data are reported as Mean (standard error).
Asterisks indicate significant differences between age groups (n.s. $p > 0.05$ ; \* $p < 0.05$ ; \*\* $p < 0.01$ ; \*\*\* $p < 0.001$ ).

### 2.5 Event-based Analysis for Autonomic/Central Event

#### 2.5.1 Time-locked analysis

In order to calculate changes in SWA and sigma power around the HR burst, the Hilbert transform was applied on filtered EEG signals in bands of interest (0.5–1 Hz for SWA and 12–15 Hz for sigma activity). To assess the HF amplitude fluctuation around the HR burst (RR_HF_), the Hilbert transform was applied on RR_HF_ (0.15–0.4 Hz). See Naji et al 2019 for detailed methods.

#### 2.5.2 ACE Change scores

We investigated ACE coupling during wake and sleep stages by tracking fluctuations in the EEG in a 20-sec window from 10 second before to 10 second after the HR burst peak. As we were specifically interested in sleep EEG activity previously demonstrated to correlate with cognitive enhancement, we focused on SWA and sigma activity. In addition, we examined RR_HF_ in the ECG channel, about which we did not have a specific hypothesis.

EEG/ECG data were binned into 5-sec intervals within the 20-sec windows around the HR burst, named -10, -5, +5, +10 window. For ACE activity, the average RR_HF_, SWA, and sigma activity were calculated in each of the four 5-sec windows. For non-ACE brain activity, we calculate average RR_HF_, SWA, and sigma activity in periods with no HR burst (including 20 s windows around them). We computed ACE change scores for each 5-sec interval as follows: (ACE activity in each 5 s interval – non-ACE activity)/ (ACE activity in each 5 s interval + non-ACE activity). Figure 1 showed an example of ACE time-locked analysis. Summary for ACE change scores were shown in Table 5 and Figure 2.

**Figure 1.**
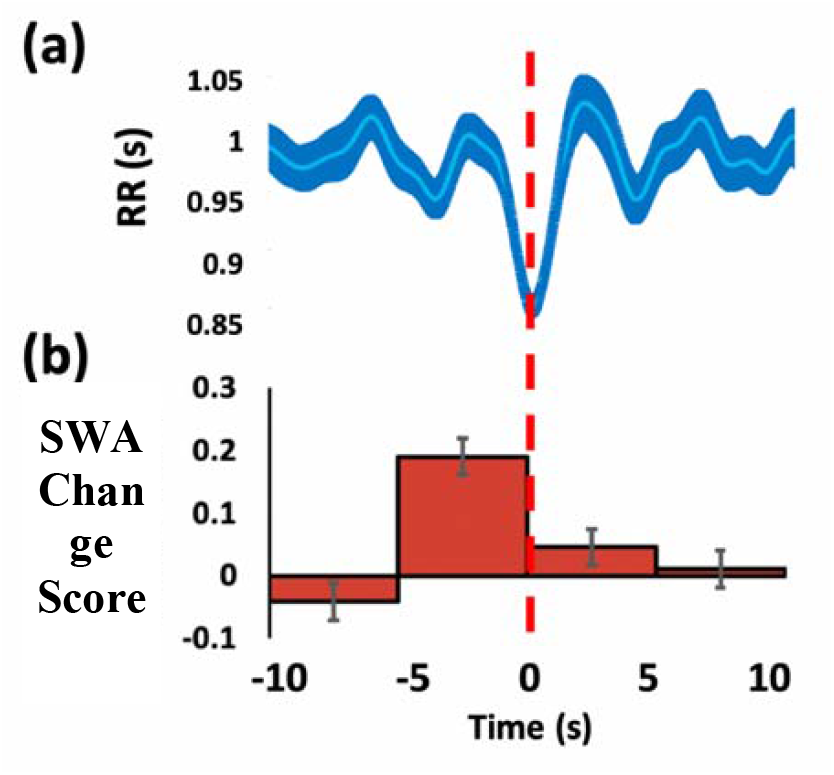
Illustration of time-locked analysis: Heart-rate-bursts and SWA change scores time-locked on HR bursts. (a) We first identified HR bursts and time-locked on the peak of HR burst to calculate EEG power spectrum within each 5-sec window. (b) Next, we averaged EEG power spectrum within individuals and calculated a change score for each individual as follows: (ACE activity in each 5 s interval – non-ACE activity)/ (ACE activity in each 5 s interval + non-ACE activity).

**Table 5.1:** Change Scores for ACE variables during N2. Data were shown in mean (standard error of the mean).

| Window |  | -10 | -5 | +5 | +10 |
| --- | --- | --- | --- | --- | --- |
| SWA | Young | -0.0239(0.0127) | 0.1453(0.0174) | -0.0354(0.0126) | -0.0524(0.0133) |
|  | Older | 0.0122(0.0182) | 0.0999(0.0227) | -0.0302(0.0179) | -0.0198(0.0223) |
| Sigma | Young | 0.0634(0.0260) | 0.1220(0.0252) | 0.0500(0.0223) | -0.0004(0.0165) |
|  | Older | 0.0403(0.0160) | 0.1001(0.0199) | 0.0767(0.0228) | 0.0274(0.0126) |
| RR<sub>HF</sub> | Young | 0.0058(0.0086) | 0.0622(0.0145) | 0.1794(0.0138) | 0.0288(0.0101) |
|  | Older | 0.0034(0.0284) | 0.187(0.0441) | 0.2137(0.0391) | 0.008(0.0217) |

**Table 5.2:** Change Scores for ACE variables during N3. Data were shown in mean (standard error of the mean).

| Window |  | -10 | -5 | +5 | +10 |
| --- | --- | --- | --- | --- | --- |
| SWA | Young | -0.0150(0.0141) | 0.1019(0.0184) | 0.0200(0.0159) | 0.0024(0.0153) |
|  | Older | -0.0280(0.0226) | 0.0125(0.0339) | 0.0011(0.0222) | 0.0008(0.0212) |
| Sigma | Young | -0.0020(0.0107) | 0.0355(0.0075) | -0.0008(0.0116) | -0.0096(0.0100) |
|  | Older | 0.0010(0.0125) | 0.0228(0.0154) | 0.0326(0.0152) | -0.0003(0.0132) |
| RR<sub>HF</sub> | Young | 0.0222(0.0162) | 0.0764(0.0218) | 0.1383(0.0222) | 0.0449(0.0217) |
|  | Older | -0.0261(0.0326) | 0.1741(0.0588) | 0.2470(0.0514) | 0.0062(0.0243) |

**Table 5.3:** Change Scores for ACE variables during Wake. Data were shown in mean (standard error of the mean).

| Window |  | -10 | -5 | +5 | +10 |
| --- | --- | --- | --- | --- | --- |
| SWA | Young | 0.1101(0.0394) | 0.2095(0.0414) | 0.0703(0.0444) | -0.0156(0.0399) |
|  | Older | 0.1009(0.0342) | 0.2273(0.0419) | 0.0795(0.0410) | 0.0078(0.0347) |
| Sigma | Young | 0.0634(0.0260) | 0.1220(0.0252) | 0.0500(0.0223) | -0.0004(0.0165) |
|  | Older | 0.0403(0.0160) | 0.1001(0.0199) | 0.0767(0.0228) | 0.0274(0.0126) |
| RR<sub>HF</sub> | Young | -0.0809(0.0428) | -0.0581(0.0538) | 0.0450(0.0469) | -0.0757(0.0519) |
|  | Older | -0.0207(0.0549) | 0.1912(0.0628) | 0.1817(0.0602) | -0.0680(0.0513) |

**Figure 2.**
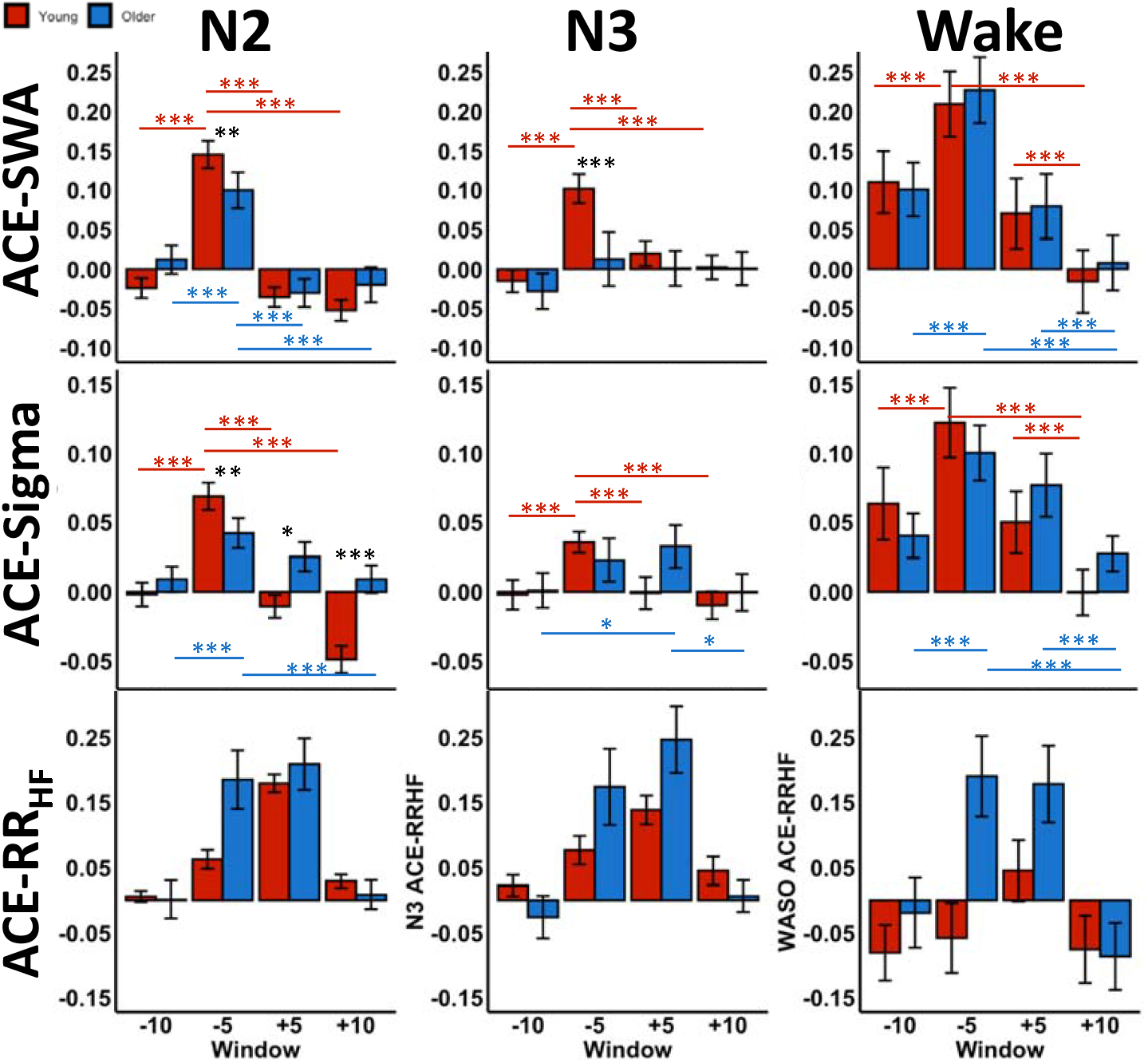
ACE change scores during the four windows across HR burst in younger (red) and older (blue) adults. Error bars represent standard error of the mean. X axis represents the four 5-sec intervals within the 20-sec windows around the HR burst, named -10, -5, +5, +10 window. Y axis represents ACE change score during the four windows. Asterisks indicate three-way post-hoc comparisons corrected by Bonferroni method. Black asterisks indicate differences between age groups within the same windows. Red asterisks indicate differences between windows within the young adults. Red asterisks indicate differences between windows within the young adults. Blue asterisks indicate differences between windows within the older adults.

**Figure 3.**
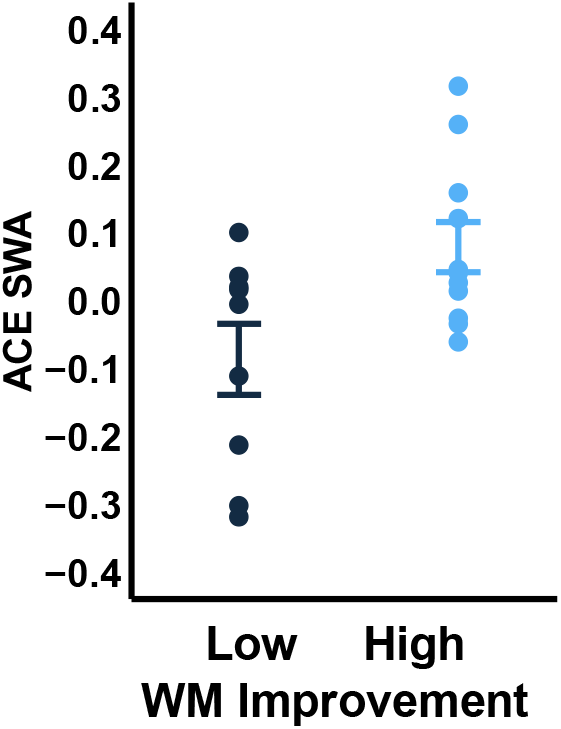
ACE-SWA during the -5 windows between older adults with high (light blue) vs low (dark blue) working memory improvement. Error bars represent standard error of the mean. Y axis represents ACE-SWA during the -5 windows.

### 2.6 Statistics Analyses

Statistical analyses were conducted using R version 3.4.3, and alpha level was set at <.05. In order to investigate within-subject profiles of ACE activity across sleep stages, channels, and windows, we used a linear-mixed effect model (LME), which does not depend on limited assumptions about variance-covariance matrix assumptions (sphericity). Additionally, LME models eliminates the need for averaging epochs across sleep stages and allows inclusion of an unbalanced number of observations per subject in the analyses. Moreover, LME models consider the influence of factors whose levels are extracted randomly from a population (i.e. participants), thus yielding more generalizable results. For RR intervals and HR bursts Density, we built an LME model using participant as crossed random effects, with a between-subject factor (two age groups) and a within-subject factor (sleep stages: N2, N3, Wake). For ACE-SWA and ACE-Sigma change scores, we built an LME model using participant as crossed random effect, with a between-subject factor (two age groups) and two within-subject factor (windows: -10, -5, +5, +10; sleep stages: N2, N3, Wake) as within-subject fixed effects. ACE-SWA and ACE-sigma change scores were averaged across frontal and central areas (F3, F4, C3, and C4 channels) in the statistical models given our preliminary analyses showing no significant topographical differences in ACE change scores. For ACE-RR_HF_ change scores, we built an LME model using participant as crossed random effect, with a between-subject factor (two age groups) and two within-subject factor (windows: -10, -5, +5, +10; sleep stages: N2, N3, Wake) as within-subject fixed effects. Post hoc comparisons were corrected using the Bonferroni method.

To examine the relationship between overall EEG power and modulation of EEG around HR burst, Pearson’s correlations were used to test the bivariate correlations between overall sigma power and ACE-Sigma change scores, as well as overall SWA power and ACE-SWA change scores.

## 3. Results

### 3.1 Sleep Architecture

Older adults demonstrated shorter total sleep time, longer N1 sleep duration, shorter N3 sleep duration, shorter REM sleep duration, longer wake after sleep onset, and lower sleep. Efficiency. Descriptive statistics for sleep architecture were shown in Table 2.

### 3.2 Age Differences in RR intervals and HRB Density (per Minute)

We first examined age differences in RR intervals (see Table 4), and found a main effect of Sleep stages (F_(2, 113)_ = 27.628, p < .0001), with heart rate slower during N2 and N3 compared to Wake (all ps < .0001); and an interaction between sleep stages and age (F_(2, 113)_ = 5.366, p = .0059) on RR intervals. No significant main effect of age was found (F_(1, 101)_ = 3.777, p = .0547). Post-hoc comparisons revealed that the young adults, compared to the older adults, showed a significant slower heart rate during N2 (p = .0164) and N3 (p = .0202) but not during Wake (p = .4288). Within each age group, the young adults showed significantly faster heart rate during Wake, compared to all other sleep stages (all ps < 0.0001), while the older adults showed no differences across sleep stages (all ps > 0.1022).

Next, we found a main effect of sleep stages (F_(2, 141)_ = 52.0966, p < .0001) and age (F_(1, 102)_ = 8.1390, p =.0052), as well as an interaction between Sleep stages and Age (F_(2, 141)_ = 15.4628, p <.0001) on HR burst Density (see Table 4). Post hoc comparisons revealed that the young adults, compared to the older adults, showed significant more HRBs during N2 (p <.0001) and N3 (p =.0002), but not during Wake (p = .3120). Within each age group, the young adults showed significantly less HRBs during Wake, compared to all other sleep stages (all ps < .0001), while the older adults significantly less HRBs during Wake, compared to N2 (p = .0028) but not N3 (p = .2211). Summary for HR bursts Density were shown in Table 4.

In summary, age-related decrease in RR intervals and HRBs density were found during NREM sleep but not Wake, suggesting a sleep-specific decline in cardiac autonomic control with aging.

### 3.3 ACE-SWA change score is modulated by ACE Window and Age

Next, we examined ACE-SWA change scores (see Figure 2) with a between-subject factor (two age groups) and two within-subject factors (windows: -10, -5, +5, +10; sleep stages: N2, N3, Wake). The analysis revealed a main effect of sleep stages (F_(2, 3389)_ = 139.7949, p <.0001), a main effect of windows (F_(3, 3389)_ = 186.9396, p<.0001), a two-way interaction between sleep stages and age groups (F_(2, 3389)_ = 5.7078, p = .0034), a two-way interaction between sleep stages and windows (F_(6, 3389)_ = 16.9731, p < .0001), a two-way interaction between age groups and windows (F_(3, 3389)_ = 6.0064, p = .0004), a three-way interaction between windows, age, and sleep stages (F_(6, 3389)_ = 2.2528, p =.0358).

The Bonferroni post-hoc comparisons revealed that during N2 and N3, young adults showed greater ACE-SWA change scores than the older adults during the -5 window (N2: p = .0068; N3: p = .0001) but no age differences between the rest of the windows (all ps > .2117). No age-related differences were found during Wake (all ps > .4379).

Within each age group, young adults showed significantly greater ACE-SWA change scores during the -5 window compared to the rest of the windows during N2 and N3 (all ps < .0001), while older adults showed significantly greater ACE-SWA change scores during the -5 window compared to the rest of the windows during N2 (all ps < .0001) but not during N3 (all ps > 0.3845). During Wake, both age groups showed significant pairwise differences between windows (all ps < .0001) except that no differences were found between the -10 and +5 windows (all ps > .2379).

Taken together, age-related decrease in peak ACE-SWA change scores were found during NREM sleep but not Wake, suggesting a sleep-specific decline in central autonomic coupling with aging. Interestingly, our older subjects showed a similar but less robust ACE modulation, with the peak EEG power during N2, and no ACE modulation during N3.

### 3.4 ACE-sigma change score is modulated by ACE Window and Age

Similarly, we examined ACE-sigma change scores (see Figure 2) with a between-subject factor (two age groups) and two within-subject factors (windows: -10, -5, +5, +10; sleep stages: N2, N3, Wake). The analysis revealed a main effect of sleep stages (F_(2, 3389)_ = 146.6889, p < .0001), a main effect of windows (F_(3, 3389)_ = 131.9417, p < .0001), a two-way interaction between sleep stages and windows (F_(6, 3389)_ = 6.7877, p < .0001), a two-way interaction between age groups and windows (F_(3, 3389)_ = 26.8665, p<.0001), a three-way interaction between Windows, Age, and Sleep stages (F_(6, 3389)_ = 2.6803, p = .0135).

The Bonferroni post-hoc comparisons revealed that during N2, young adults showed higher ACE-Sigma change scores during the -5 window than the older adults (p =.0055) but lower during the +5 (p = .0111) and +10windows (p < .0001). During N3, older adults showed higher ACE-Sigma change scores during the +5 window (p = .0492) than the young adults but not the rest of the windows (all ps > .1411). During Wake, older adults showed higher ACE-Sigma change scores during the +5 (p = .0128) and +10 window (p = .0101) than the young adults but not the rest of the windows (all ps >.1406).

Within each age group, during N3, Bonferroni post-hoc comparisons revealed significantly greater ACE-Sigma change scores during the -5 window compared to the rest of the windows (all ps < .0003) among the young adults, whereas the older adults showed significantly greater ACE-Sigma change scores during the +5 window compared to the -10 and +10 windows (all ps <.0383). During N2, young adults showed significantly greater ACE-Sigma change scores during the -5 window compared to the rest of the windows (all ps < .0001) and lower ACE-Sigma change scores during the +10 window compared to the -10 and +5 windows (all ps < .0001), whereas the older adults showed significantly greater ACE-Sigma change scores during the -5 window compared to the -10 and +10 windows (all ps < .0007). During Wake, young adults showed significantly greater ACE-Sigma change scores during the -5 window compared to the rest of the windows (all ps < .0001) and lower ACE-Sigma change scores during the +10 window compared to the -10 and +5 windows (all ps < .0001), and the older adults showed similar pattern, with significantly greater ACE-Sigma change scores during the -5 window compared to the rest of the windows (all ps < .0376) and greater ACE-Sigma change scores during the +5 window compared to the -10 and +10 windows (all ps < .0002).

Taken together, age-related decrease in peak ACE-Sigma change scores were found during N2 but not N3 or Wake. Similar to ACE-SWA, our older subjects showed a similar but less robust ACE modulation, with the peak EEG power during N2, and no ACE modulation during N3. Again, no age-related differences were found in ACE-Sigma during Wake, suggesting a sleep specific decline in heart-brain coupling.

### 3.5 ACE-RR_HF_ change score is modulated by ACE Window and Age

In addition to EEG fluctuation, we examined ACE-RR_HF_ change scores (see Figure 2) with a between-subject factor (two age groups) and two within-subject factors (windows: -10, -5, +5, +10; sleep stages: N2, N3, Wake). The analysis revealed a main effect of sleep stages (F_(2, 984)_ = 7.5304, p = .0006), a main effect of windows (F_(3, 984)_ = 60.5159, p < .0001), a two-way interaction between sleep stages and age groups (F_(2, 984)_ = 5.7119, p =.0034), a two-way interaction between age groups and windows (F_(3, 984)_ = 10.8937, p < .0001). No significant three-way interaction between age groups, sleep stages, and windows was found (F_(6, 984)_ = 0.7177, p = .6354).

The Bonferroni post-hoc comparisons revealed that regardless of sleep stages, older adults showed higher ACE-RR_HF_ change scores during the -5 window than the young adults (p = .0017) but not during the rest of the windows (p > .0795). Within each age groups, young adults showed significantly greater ACE-RR_HF_ change scores during the +5 window than the rest of the windows (all ps < .0023), whereas the older counterparts showed significantly greater ACE-RR_HF_ change scores during both the -5 and +5 window than the rest two windows (all ps < .0001). In addition, regardless of windows, young adults showed greater ACE-RR_HF_ change scores during N2 and N3 compared to Wake (all ps < .0002), whereas the older adults showed no significant differences between sleep stages (all ps > .9307).

In summary, young adults demonstrated peak RR_HF_ during the +5 window, similar to Naji et al. (2019)a. However, the older counterparts showed an earlier rise of RR_HF_ with the peak amplitude during both the -5 and +5 windows.

### 3.6 Relationship between Overall EEG Power and ACE EEG Change Scores

One possibility is that the overall EEG power loss in older adults contributed to the individual differences in ACEs declines. As such, we examined overall EEG power across the whole sleep period regardless of ACE windows in the SWA and Sigma bands, and confirmed an age-related loss in EEG power, with older adults demonstrating lower SWA power during N2, N3 and Wake, as well as less sigma power during N3 sleep. Summary statistics for overall EEG power during each sleep stage were shown in Table 3.

Next, we used Pearson correlation coefficients to examine the bivariate relationship between overall EEG power and ACE EEG change scores. No significant associations between overall SWA power and ACE-SWA change score during N2 (Young: all ps > .3564; Older: all ps > .0956), N3 (Young: all ps > .1783; Older: all ps > .5319), or Wake (Young: all ps > .1540; Older: all ps > .2972). Similarly, no significant associations between overall Sigma power and ACE-Sigma change score during N2 (Young: all ps > .3239; Older: all ps > .3184), N3 (Young: all ps > .1861; Older: all ps > .4239), or Wake (Young: all ps > .3575; Older: all ps > .0880). Taken together, these suggested that the age-related differences in EEG modulation around HR bursts cannot be explained by overall EEG loss in older adults.

### 3.5 ACE-SWA during N3 Associated with Working Memory Improvement in Older Adults

To extended our previous findings of an association between ACE and working memory improvement in young adults, we tested the relationship between ACE-SWA change scores during N3 and WM improvement measured with the Operation-Span task among our older subjects. Since only 23 older participants had stable N3 sleep for ACE analysis, we computed a median-split of their WM improvement scores and compared ACE-SWA change scores in the two groups using a non-parametric Wilcoxon signed-rank test. We found that people who showed greater WM across the nap were also the people who had larger ACE-SWA change scores during N3 (F3: p = 0.0232; F4: p = 0.0877; C3: p = 0.0402; C4: p = 0.3832). These data suggest that, similar to younger adults, high-functioning WM processing may reflect greater coupling between the autonomic and central nervous systems, whereas poor coupling suggests potential impairment to the prefrontal network supporting WM.

## 4. Discussion

We aimed to characterize the impact of age on autonomic-central couplings profiles across wake and sleep, and the functional significance of this coupling for working memory improvement across sleep in older adults. To this end, we compared younger and older adults in Autonomic-Central Events (ACEs) during daytime naps. In the young adult population, our previous sleep findings were replicated in the present study: During wake, N2 and N3, a significant increase in SWA and sigma activity preceded the HRB peak. In contrast, our older subjects demonstrated lower amplitude of ACE modulation during the -5 window in both N2 and N3, but not wake. Replicating our original study, we also observed in younger adults a significant vagal response with increased RR_HF_ after the HRB peak. In contrast, for older adults, we observed a reduced amplitude of ACE-SWA and ACE-Sigma during NREM sleep, and an earlier-onset and sustained RR_HF_ rise before and after the HRB peak. Furthermore, we found that the change in age-related ACEs was not associated with changes EEG sleep features. Interestingly, using a median-split analysis based on WM performance we showed that greater greater WM improvement across the nap was associated with more robust ACE-SWA during N3 in older adults, Altogether, our results reflect that with age, subjects show a decline in temporal coupling between the autonomic and central nervous systems, and that this decline reflects individual differences in WM improvement.

### Age-Related Declines in Cardiac Autonomic Control

Reduced cardiac autonomic control is a typical consequence of growing older. Possible influential factors include reduced SA node cell functionality, which results in greater time between heart beats; lower maximal aerobic exercise capacity caused by reduced physiological functioning; and reduced α and β adrenergic receptor sensitivity in the heart as a result of increased sympathetic activity (Hagberg et al., 1985; Hawkins & Wiswell, 2003; Jones Pamela Parker et al., 2003; Rowe & Troen, 1980; Zhang, 2007). Consistent with prior research, we found that older adults had faster HR than younger adults during NREM sleep. However, older adults’ rate of HRBs during sleep was half that observed in younger adults during NREM but not Wake, with young adults having approximately one HRB per minute while older adults had one approximately every two minutes. This loss of sleep-associated HRBs in older adults may be the result of age-associated heart rate changes or could reflect reduced vagal cardiac control, as faster baseline heart rates and lower cardiac controls might not provide the best state for periodic heart rate acceleration. In line with our findings, de Zambotti et al. (2016) showed that tone-evoked KCs that followed by heart rate acceleration were more likely to be elicited when the preceding heart rates were slower, suggesting that specific ANS states may facilitate cortical synchronization (de Zambotti et al., 2016; Otzenberger et al., 1997; Trinder, 2007). Typically, the vagus nerve reduces the heart rate via a negative feedback loop, termed the vagal break, in which increased vagal activity reduces the heart rate on a beat-to-beat scale [58]. By observing the age-dependent loss of HRBs, we are likely seeing a less effective vagal break in our older adult subjects.

### Autonomic-Central Coupling: Mechanisms and Implications with Aging

Prior literature has suggested that autonomic activity may be linked with sleep brain activity, and that this interaction is likely a distinct predictor of plasticity, cognitive ability and enhancement [10,40,41,50]. Studies have revealed a consistent symmetry between heart and brain activity with temporal changes in NREM delta (0.5-4Hz) power, a marker of homeostatic sleep drive, and ANS activity (Ako et al., 2003; Brandenberger et al., 2001; Jurysta et al., 2003, 2005; Kuo & Yang, 2004; Rothenberger et al., 2015; Thomas et al., 2014; Yang et al., 2002). In addition to macro-structure, at the micro-structure level, de Zambotti et al. (2016) found that both auditorily evoked and spontaneous KCs were quickly followed by brief increases in heart rate. Similarly, Naji et. al (2019) and Chen et al. (2020) demonstrated a temporally coincident increase in SWA and Sigma preceding HRBs in NREM sleep. In the current study, we replicated our ACE observations in young adults, and found a sleep-specific decline in such coupling among older adults.

The driving force behind the age-related decline of ACEs is unclear, and the ability to identify that force is further complicated by a lack of consensus regarding communication flow between heart and brain. While mechanisms driving such autonomic-central couplings remains unclear, some evidence points to arousal response or the gating mechanisms of sleep. If the autonomic activation (tachycardia, or HR bursts) is a sign of arousal (sleep discontinuity), the finding of KCs and SWA during enhancement of autonomic activation indicates the possibility of physiological activation without sleep disruption. Initially the cortex tries to preserve sleep continuity with reinforcement of its gates (KCs and SWA; sleep-protective response). When the thalamic gate cannot control the afferent inputs, a cortical arousal (alpha/beta frequency burst) might be observed. In the current study, boost of SWA and Sigma activities might indicate the amplitude of the adaptive sleep-protective process, and therefore, is associated with successful aging indicated by greater working memory functions. In line with our findings, de Zambotti et al. (2016) showed that tone-triggered KCs are temporally coupled with a rapid increase and then decrease in heart rate activity, and coincide with bursts of KCs and slow waves (Sforza et al., 2000). In this context, both synchronous EEG (KCs, or bursts of SOs) and cardiovascular activations (heart-rate acceleration) were viewed as responses to arousal from sleep, as when acoustic tones were not accompanied by KCs, the heart rate fluctuation was reduced or absent, indicating that arousal responses might be driving ANS/CNS activities. It’s been hypothesized that in the case of the KCs, the recruited synchronized EEG response acts as a mechanism to decrease cortical arousal, suggesting that the heart-rate acceleration (tachycardia, or HR bursts) can be viewed as peripheral response of arousal from sleep. Taken together, the subsequent heart-rate deceleration that de Zambotti et al. (2016) showed and the surge of HF that Naji et al. (2019) and the current study found may reflect a feedback effect of arousal showing an inertial effect once the arousal stimulus is removed. Alternatively, arousal and post-arousal periods may modulate the autonomic system reflecting the activation-deactivation of neuronal oscillations intrinsically regulated by the cyclic arousability of the sleepy brain (Cyclic Alternating Pattern; Schnall et al., 1999; Sforza et al., 1999; Ferri et al., 2000). Interestingly, the age-related ACEs decline we observed parallel the previous finding of a decreased rate of subtype A1 Cyclic Alternating Pattern (KCs and delta bursts co-occur with autonomic activation) with aging. These age-related decline in ACEs might reflect a dysfunctional gating mechanism which results to instability of sleep.

In addition to arousal responses, ACEs may also represent the top-down control on brainstem’s dynamic maintenance of homeostasis during the transition from wake to sleep. As homeostatic pressure drives the transition from lighter sleep to deeper stages, the CNS and ANS experience large and rapid slowing in physiological rhythms, including decreased heart rate, broad synchronization of EEG slow waves (Fernandez Guerrero & Achermann, 2018), as well as alignment of cortical and autonomic signals (Ulke et al., 2017). The neurovisceral integration model may provide insight into the directionality of the heart-brain interplay pathways, at it has implicated several candidate brain structure networks in the communication and regulation of cognitive, emotional, and autonomic function (Thayer & Lane, 2000). Brainstem medullary nuclei are responsible for a wide range of bodily functions including deepening of slow wave sleep (Anaclet & Fuller, 2017) and deceleration of heart rate (Monge Argilés et al., 2000). Baroreceptors in the carotid sinus and aortic arch send messages to the vagal nerve which are relayed to the nucleus of solitary tract (NTS) in the medulla, which further projects the message to higher-order cognitive areas such as hippocampus, amygdala, and prefrontal cortex, which can initiate a top-down inhibitory mechanism that initiate parasympathetic control of the heart and slow it down (Silvani et al., 2011; van der Kooy et al., 1984). Increased vagal activity can subsequently activate the projecting NTS pathways to the basal forebrain and locus coeruleus, which can result in an increased release of acetylcholine and norepinephrine leading to a reduction of slow oscillations (Krishnan et al., 2016). It is, therefore, possible that medullary nuclei regulating wake to sleep transitions modulate the pace of the slow down with brief accelerations in cardiac activity. Due to the relative overlap between nuclei, these fluctuations may promote sleep promoting oscillations such as SOs and spindles coupled with heart rate accelerations. Thus, ACEs may represent the adaptive and flexible modulation of central and peripheral activities by the brainstem. Therefore, the decline of ACEs might represent a reduced top-down prefrontal controls on subcortical and peripheral regions through the brainstem. Such reduced prefrontal functioning is also implicated in the individual differences in working memory in our other subjects. However, more research is needed to make definite conclusions.

Altogether, while the directionality of communication pathways between the heart and brain is not definitive, our findings of ACEs and its declines are in line with the previous literature. Overall age-related neural degeneration, primarily in frontal cortex, may cause impaired pathway communication, resulting in less successful control of ANS activity as well as poorer temporal convergence of the two nervous system branches.

### Autonomic-Central Coupling and Cognitive Decline

Age-related cognitive decline is a major public health concern as the global population rapidly ages. In particular, deficits in working memory play a central role in normal neurocognitive aging and the rapid cognitive deterioration associated with dementia, including Alzheimer’s disease. The underlying cause of age differences in working memory has been hypothesized to involve alterations in the functional connectivity of large-scale brain networks that normally function in a coordinated or synchronous fashion (Tomasi & Volkow, 2012). During sleep, temporal alignment between thalamic sleep spindles and the up-state of cortical slow oscillations has been implicated in sleep-dependent cognitive enhancement (Latchoumane et al., 2017), and decreased coupling among older adults have been implicated in decreased sleep-dependent benefits in both long-term (Helfrich et al., 2019) and working memory decline (Sattari et al., 2019). Here, we demonstrated that older adults with a greater sleep-dependent working memory improvement also showed a greater ANS-CNS coupling, a finding aligned with our young subjects (Chen et al., 2020). Synchronization between ANS and CNS may reflect prefrontal cortex – subcortical inhibitory control processes supporting working memory. In older adults with mild cognitive impairment undergoing a six-week training intervention, Lin et al. (2017) revealed a link between autonomic vagal activity and cognitive improvements measured across a range of executive functions including working memory (Lin et al., 2017). They further showed that cognitive training enhanced HF-HRV and decreased connectivity between striatal and prefrontal regions, suggesting that vagal activity might reflect enhanced cognitive control via greater automaticity and reduced activation between the striatum and prefrontal networks (Lewis et al., 2004). Although sleep was not measured across the cognitive training intervention, the current findings suggest that the strengthening of prefrontal-vagal networks supporting performance improvement may have occurred during sleep.

Further research is needed to determine the causal role and the neurological mechanisms of ACE on age-related cognitive decline. Research efforts in this direction will lay the groundwork for developing future non-invasive interventions aimed at reducing cognitive deficits in physiological aging and clinical populations.

## Author contributions

S.C.M. conceived and designed research; L.N.W. and N.S. collected data and participant responses; P.C. analyzed data and prepared figures; P.C. drafted the introduction, methods, results, and discussion; K.C.S drafted the introduction; P.C. and S.C.M edited and revised manuscript.

