## Supplemental Figures and Tables for "Autonomic/Central Coupling during Daytime Sleep Differs between Older and Younger People"

**Age-related differences in Autonomic/Central Coupling during a Daytime Nap**

**Age Group is Associated with ACE Misalignment**

Upon visual examination (Supplemental Figure S1-3), young adults demonstrated a uniform pattern with peak EEG occurring during the -5 window, whereas a less clear picture emerged in older adult group with some reaching the peak EEG during the +5 window. Similarly, for RR_HF_, most of the young adults reached the peak RR_HF_ during the +5 window, whereas some older adults reached the peak EEG during the -5 window and decreased during the +5 window. Interestingly, older individuals who showed misalignment in one measure (e.g. ACE-SWA), also more likely to have misalignment in another measure (e.g. ACE-Sigma). We devised a measure of central/autonomic misalignment by identifying if the peak ACE-EEG occurred during the +5 window or if the peak RR_HF_ occurred during the -5 window on average during a sleep stage. We then statistically examined if the number of misalignments (see Supplemental Table S1 and S2 for contingency tables; If a subject showed misalignment in ACE-Sigma and ACE-SWA during N2, this counted as misalignment = 2.) is associated with either age group by a Chi-squared independent test, and showed that being in the older adult group was associated with more misalignments (N2: chi-squared = 29.73298, p < .0001; N3: chi-squared = 13.80196; p = .0010).

Next, using a logistic regression, we investigated if the number of misalignments can be predicted by napping habits among the older adults. We found that those who reported having napping habits have a lower likelihood of showing more than one measure of misalignment during N2 (β = -1.7918, p = .0434), and that nap frequency was inversely related to the number of misalignments during N2 (β = -.4920, p = .0549). Napping habit was not correlated with the number of misalignments in N3 sleep (all ps >. 0.3749). Furthermore, the number of misalignments during either N2 or N3 cannot be predicted by STOP-Bang score, BMI, ESS, or TELE score (all ps > 0.0824). Taken together, our results suggested that central/ autonomic misalignment increased with age, and can potentially be mitigated by frequent napping.

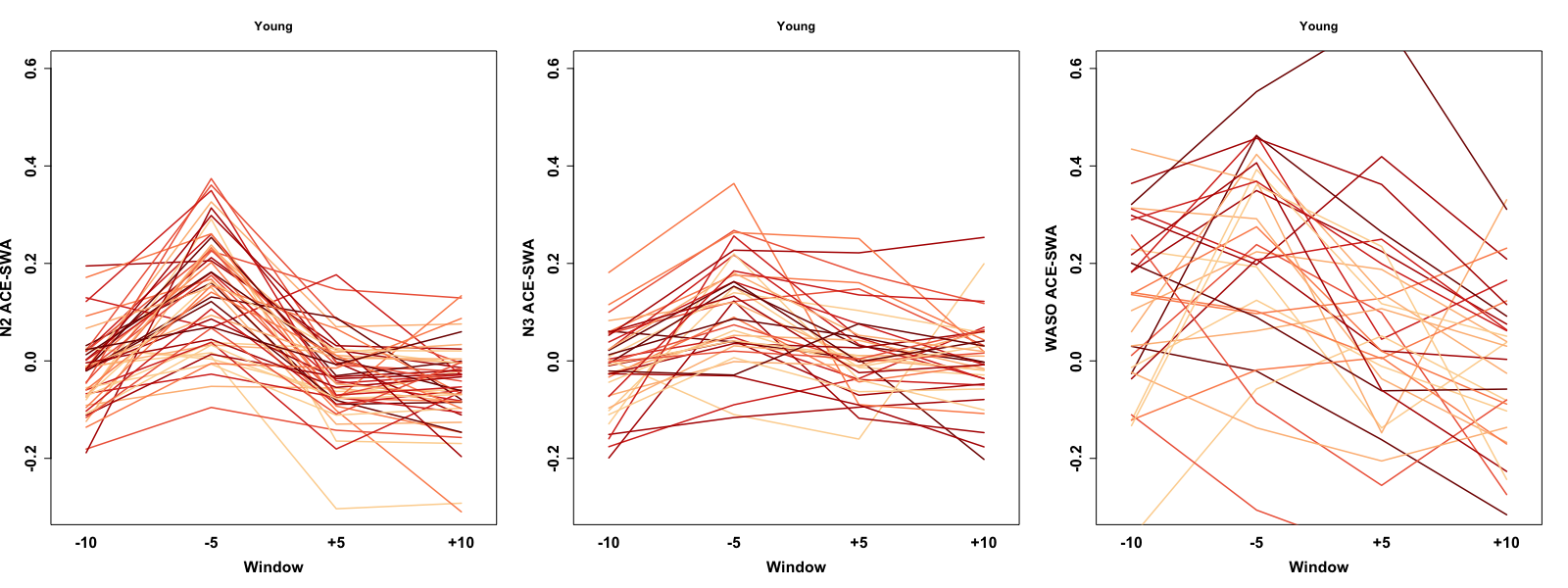

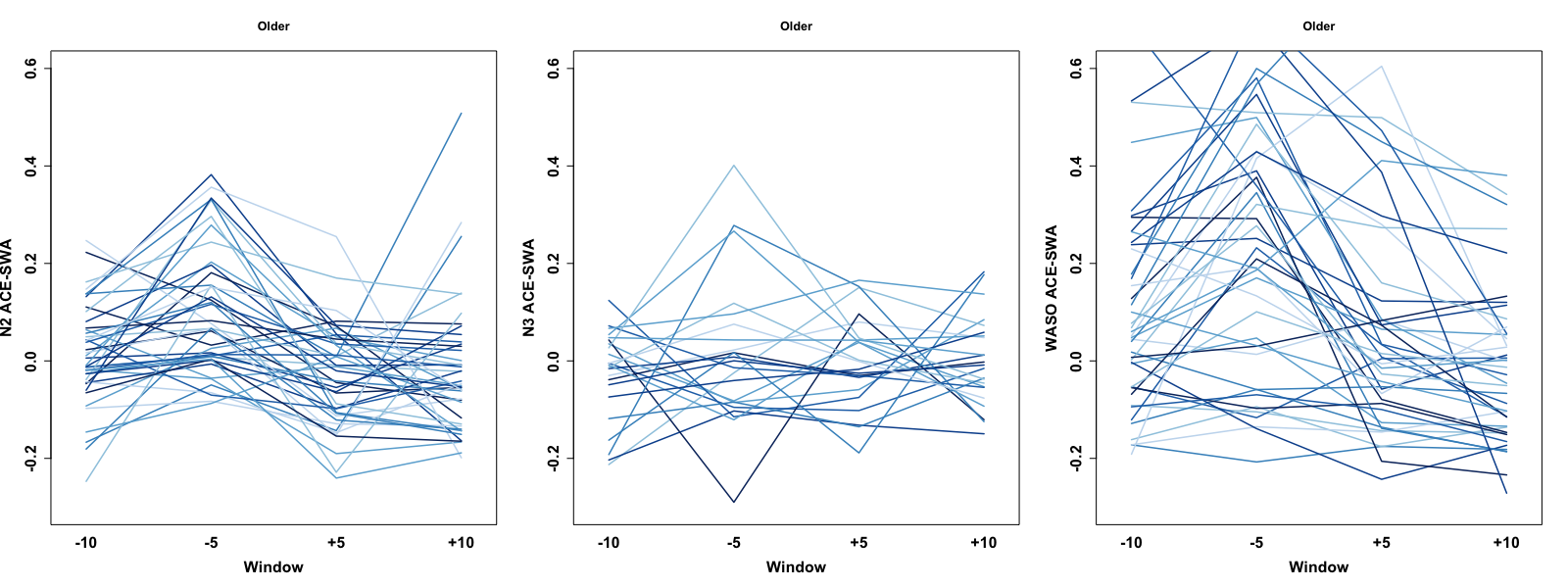

***Supplemental Figure S1***

*Individual trajectories of ACE-SWA change scores during the four windows across HR burst in younger (red) and older (blue) adults. X axis represents the four 5-sec intervals within the 20-sec windows around the HR burst, named -10, -5, +5, +10 window. Y axis represents ACE change score during the four windows.*

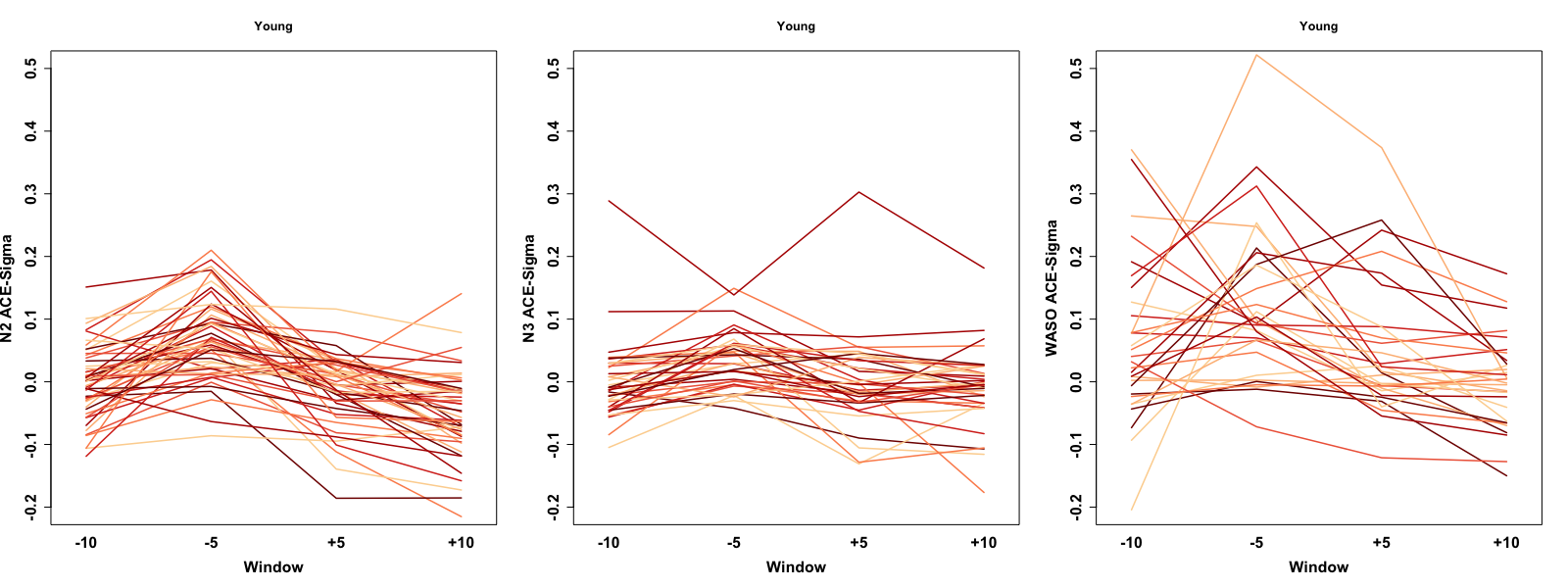

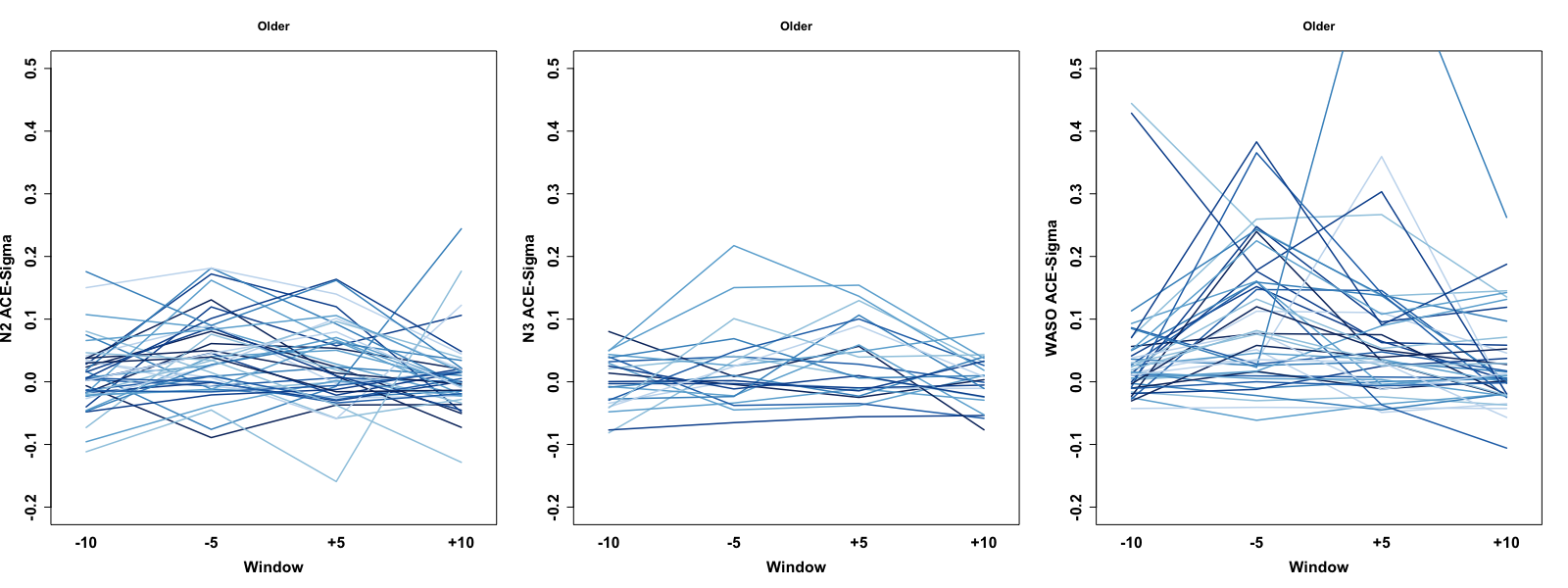

***Supplemental Figure S2***

*Individual trajectories of ACE-Sigma change scores during the four windows across HR burst in younger (red) and older (blue) adults. X axis represents the four 5-sec intervals within the 20-sec windows around the HR burst, named -10, -5, +5, +10 window. Y axis represents ACE change score during the four windows.*

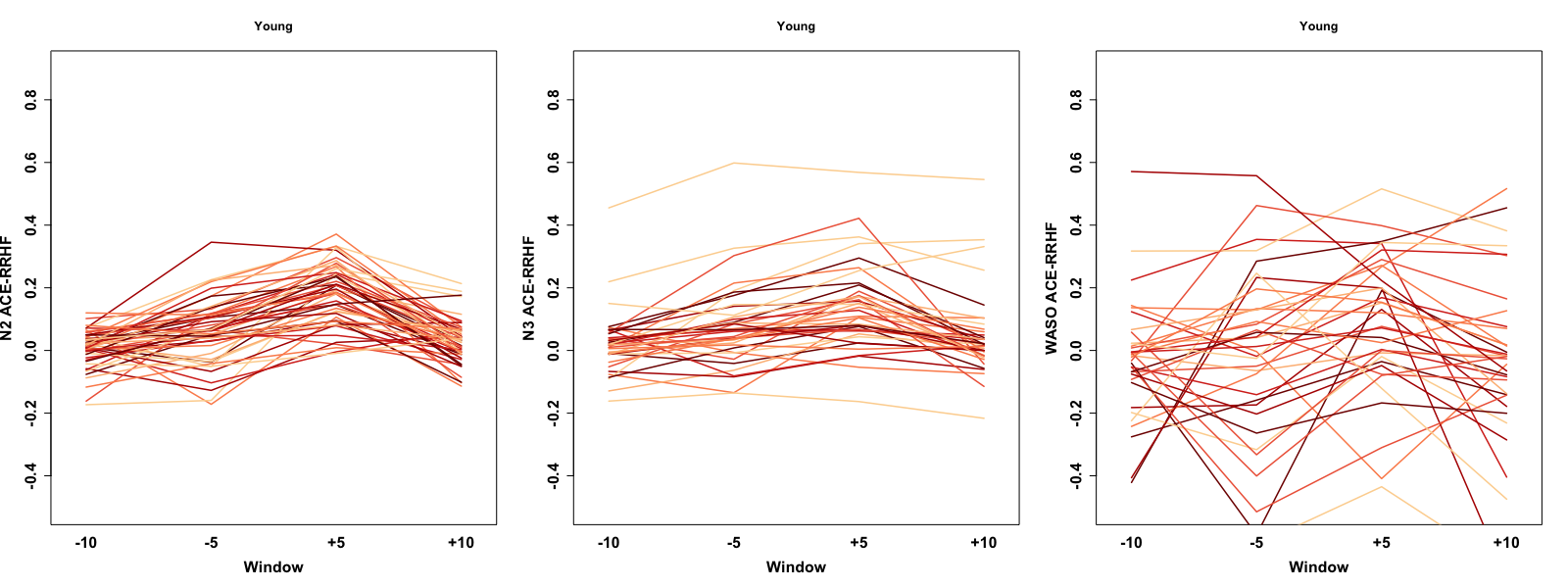

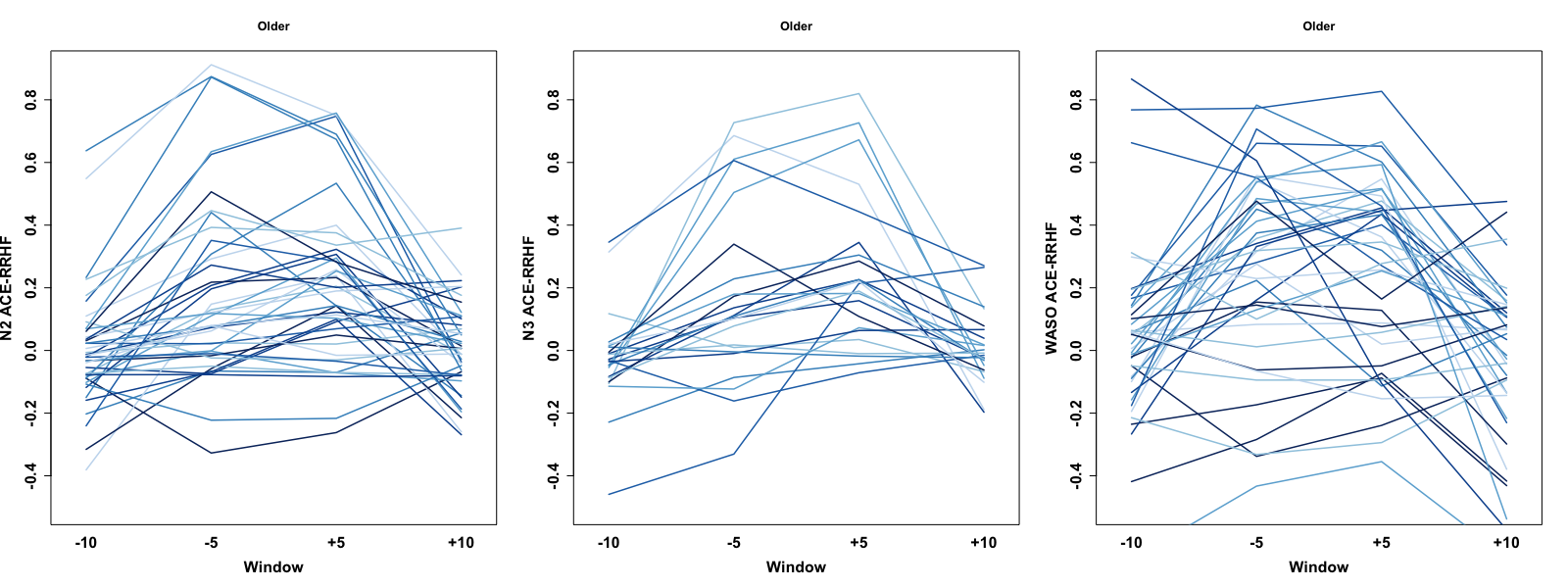

***Supplemental Figure S3***

*Individual trajectories of ACE-RR_HF_ change scores during the four windows across HR burst in younger (red) and older (blue) adults. X axis represents the four 5-sec intervals within the 20-sec windows around the HR burst, named -10, -5, +5, +10 window. Y axis represents ACE change score during the four windows.*

*Table S1 Contingency Table for Misalignment Count during N2 (chi-squared=29.73298; p< 0.0001)*

|  | | Misalignment = 0 | Misalignment = 1 | Misalignment > 1 | Total |
| --- | --- | --- | --- | --- | --- |
| Older  Younger |  | 15 | 14 | 13 | 42 |
| Younger |  | 44 | 4 | 1 | 49 |
| Total |  | 59 | 18 | 14 | 91 |

*Table S2 Contingency Table for Misalignment Count during N3 (chi-squared=13.80196; p=0.0010)*

|  | | Misalignment = 0 | Misalignment = 1 | Misalignment > 1 | Total |
| --- | --- | --- | --- | --- | --- |
| Older  Younger |  | 7 | 6 | 10 | 23 |
| Younger |  | 24 | 12 | 2 | 38 |
| Total |  | 31 | 16 | 12 | 61 |
